# Cholesterol in membranes facilitates aggregation of amyloid β protein at physiologically low concentrations

**DOI:** 10.1101/2020.09.06.285312

**Authors:** Siddhartha Banerjee, Mohtadin Hashemi, Karen Zagorski, Yuri L. Lyubchenko

## Abstract

The formation of amyloid β (1-42) (Aβ42) oligomers is considered to be a critical step in the development of Alzheimer’s disease (AD). However, the mechanism underlying this process at physiologically low concentrations of Aβ42 remains unclear. We have previously shown that oligomers assemble at such low Aβ42 monomer concentrations *in vitro* on phospholipid membranes. We hypothesized that membrane composition is the factor controlling the aggregation process. Accumulation of cholesterol in membranes is associated with AD development, suggesting that insertion of cholesterol into membranes may initiate the Aβ42 aggregation, regardless of a low monomer concentration. We used atomic force microscopy (AFM) to directly visualize the aggregation process of Aβ42 on the surface of a lipid bilayer containing cholesterol. Time-lapse AFM imaging unambiguously demonstrates that cholesterol in the lipid bilayer significantly enhances the aggregation process of Aβ42 at nanomolar monomer concentration. Quantitative analysis of the AFM data shows that both the number of Aβ42 oligomers and their sizes grow when cholesterol is present. Importantly, the aggregation process is dynamic, so the aggregates assembled on the membrane can dissociate from the bilayer surface into the bulk solution. Computational modeling demonstrated that the lipid bilayer containing cholesterol had an elevated affinity to Aβ42. Moreover, monomers adopted the aggregation-prone conformations present in amyloid fibrils. The low energy barriers between these conformations facilitate the transition between monomer states and is another factor promoting the self-association of the monomers.

## Introduction

The self-assembly of amyloid β (Aβ) proteins into aggregates is considered to be the major pathogenic event leading to the development of Alzheimer’s disease (AD) [1–3]. Evidence strongly suggests that even the smallest aggregate, a dimer, is neurotoxic and can potentially impair synapse structure and function[4, 5]. Several studies have characterized the kinetics of the Aβ aggregation process and the different factors that influence it, such as protein monomer concentration, pH of the medium, role of metal ions, and the presence of small molecules [6, 7]. However, none of these studies have been able to address the exact agent nor the mechanism of disease development; primarily because these *in vitro* studies were carried out at high micromolar concentrations, whereas the physiological concentration of Aβ is in picomolar to the low nanomolar range [8, 9]. Recently, we found that spontaneous formation of amyloid aggregates at nanomolar concentrations does occur at surfaces that play the role of catalysts.[10–12]. Experimental results, along with the theoretical modelling, demonstrated that the transient interactions of the protein monomer increase the local protein concentration, allowing for the on-surface aggregation process[13]. Importantly, biologically relevant surfaces such as phospholipid bilayers also exhibit these aggregation enhancing properties [11, 12]. Our recent study showed that Aβ42, at low concentration, can form aggregates on supported lipid bilayers composed of phospholipids 1-palmitoyl-2-oleoyl-sn-glycero-3-phosphocholine (POPC) and 1-palmitoyl-2-oleoyl-sn-glycero-3-phospho-L-serine (POPS) [12]. The on-membrane aggregation takes place at ambient conditions, physiological pH values, and with no agitation. Computer modeling revealed that the interactions of Aβ42 with the membrane significantly changed the protein structure and the altered protein conformation facilitated the aggregation process. These data are in line with a number of publications reviewed in [14] and specifically paper [15] that reported catalytic properties of zwitterionic lipid vesicles during the formation of Aβ42 fibrils. Importantly, our data showed that the on-surface aggregation is a dynamic process, and the assembled aggregate can dissociate from the surface into the bulk solution [11, 12]. Therefore, these findings suggest that the on-surface aggregation is the mechanism by which amyloid oligomers, or their seeds for aggregation, are produced. Inside the cell, the surface-assembled oligomers can induce neurotoxic effects, such as phosphorylation of the tau protein, to initiate its misfolding and aggregation.

Various factors affect the amyloid aggregation process, and lipids are among them. According to [16], the main membrane lipids present in the fibrils are cholesterol, sphingomyelin, and glycosphingolipids, such as gangliosides. Importantly, the role of the membrane lipid composition on pathogenesis of AD and facilitated assembly of amyloid fibrils is widely discussed (*e.g*., reviews [17–20]). Note a recent publication[21] in which the role of cholesterol in AD pathogenesis has been revealed. It is evident from these studies that the presence of cholesterol and gangliosides in membranes affects structure and stability of amyloid agregates and contributes to the neurotoxic effect [19], [20]. We hypothesize that the membrane composition contributes to the on-membrane aggregation process of amyloids. This hypothesis is in line with our recent data in which we demonstrated that on-surface aggregation of α-synuclein is faster on POPS bilayer compared to a POPC one [11].

Here we directly test this hypothesis and specifically focus on the role of cholesterol, incorporated in the phospholipid bilayer, on the aggregation of Aβ42 protein. We applied time-lapse atomic force microscopy (AFM) to directly monitor the aggregation of Aβ42 on POPC-POPS lipid bilayer with and without cholesterol. Our data demonstrate that cholesterol-containing bilayers show significantly higher aggregation propensity, with the faster initial appearance of aggregates and larger aggregate sizes compared to non-cholesterol bilayers. All-atom molecular dynamics simulations show that the cholesterol-containing bilayer increases the affinity of Aβ42 monomers to the bilayer surface. Moreover, there is a dramatic change of the Aβ42 conformation and a significantly greater β-structure content for the dimer in the presence of a cholesterol bilayer. We posit that these two major factors explain the catalytic effect of cholesterol in the aggregation of Aβ42 on the cholesterol-containing membranes.

## Results

### 1. Experimental studies Cholesterol accelerates the on-membrane aggregation of Aβ42

To identify the role of cholesterol on the aggregation of Aβ42, two lipid bilayers were assembled on mica substrate: 1) POPC-POPS (1:1 mol), and cholesterol (20 mol%) (PC-PS-Chol); and 2) POPC-POPS (1:1 mol) without cholesterol (PC-PS). For unambiguous identification of the Aβ42 oligomers, the lipid bilayer was prepared in such a way that it remains smooth, homogeneous, and devoid of any trapped vesicles [22]. AFM topographic images of the lipid bilayer from two different areas are shown in Fig. S1 a and b. To confirm that a bilayer has been formed on the mica substrate, the thickness of the phospholipid was measured. The height value of 4.9 nm verifies the assembly of the lipid bilayer (Fig. S1c). The supported lipid bilayer is not a static system, since the lipids are not covalently attached to the mica substrate. Therefore, its stability was monitored by time-lapse AFM imaging for 6 hr (Fig. S2). Figs. S2 a-b shows the same area of the lipid bilayer at 1 hr and 5 hr. After 6 hr of time-lapse imaging, a zoomed area was imaged (Fig. S2c); the bilayer remains smooth and does not show any defects. The roughness of the bilayer surface was measured at 1hr and 5hr, Fig. S2d, which shows no significant difference, indicating minimal change in the bilayer during the course of the experiment.

To monitor the Aβ42 aggregation on the bilayer surface, 10 nM Aβ42 monomer solution was placed on the bilayers and time-lapse AFM imaging was performed immediately after the addition; the data is shown in Fig. 1. The first frame (Fig. 1a) depicts the bilayer surface immediately after addition of the protein solution. It shows the topography of the surface with no visible aggregates. Aggregates appear after 1 hr in presence of the bilayer (Fig. 1b) and increase in size after 2 hr (Fig. 1c). Significantly larger aggregates were observed on the membrane surface after 5 hr (Fig. 1d). To quantitatively follow the aggregation process, the number and volume of the aggregates at each time point, in a 2 μm × 2 μm area, were measured and plotted in Fig. 1e. Both the number and sizes (the volume) increase monotonically over time, indicating the ongoing aggregation of Aβ42 on the bilayer surface. The volume distribution of the aggregates formed at different time points on the PC-PS-Chol lipid bilayer surface is shown in Fig. S3. The mean value of volume is obtained from the peak of the distribution. As time progresses, the mean value of the aggregate volume increases and the distribution becomes broad, sometimes with multiple peaks, indicating the formation of aggregates of widely different sizes (Fig. S3).

**Figure 1.**
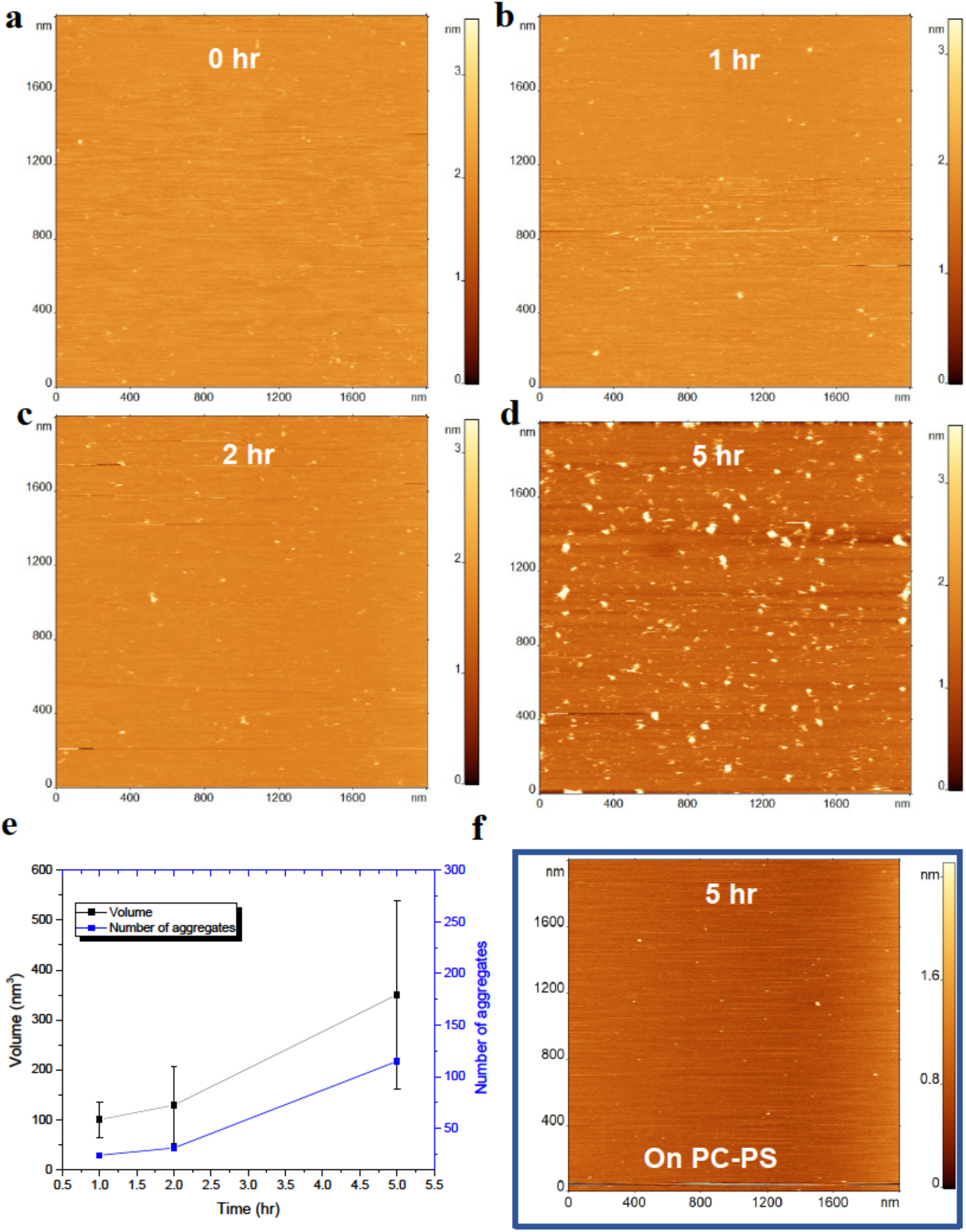
Aggregation of 10 nM Aβ42 onto PC-PS-Chol lipid bilayer. AFM topographic images of lipid bilayer surface (a) just after addition of the protein, (b) after 1 hr, (c) 2hr and (d) 5 hr. Initially the surface remains clean and as time progresses more aggregates are visible, (e) Combined plot of number and volume of the aggregates. Monotonic increase in both the number and volume is observed indicating the on-going aggregation process, (f) AFM image of the lipid bilayer after incubating 10 nM of Aβ42 for 5 hr without cholesterol on PC-PS lipid bilayer.

The Aβ42 aggregates after 5 hr on the bilayer without cholesterol is shown in Fig. 1f. The analysis shown in Figs. 2 a, b demonstrates that the aggregates present on the surface are significantly fewer and smaller compared to the PC-PS-Chol bilayer (Fig. 1).The bar plot in Fig. 2a shows the comparison of the sizes of the aggregates after 5 hr of time-lapse experiment in presence (grey bar) and absence (black bar) of cholesterol in the lipid bilayer. Furthermore, the number of aggregates which are formed on the PC-PS-Chol bilayer is approximately 6 times greater compared to the PC-PS lipid bilayer after 5 hr (Fig. 2b).

**Figure 2.**
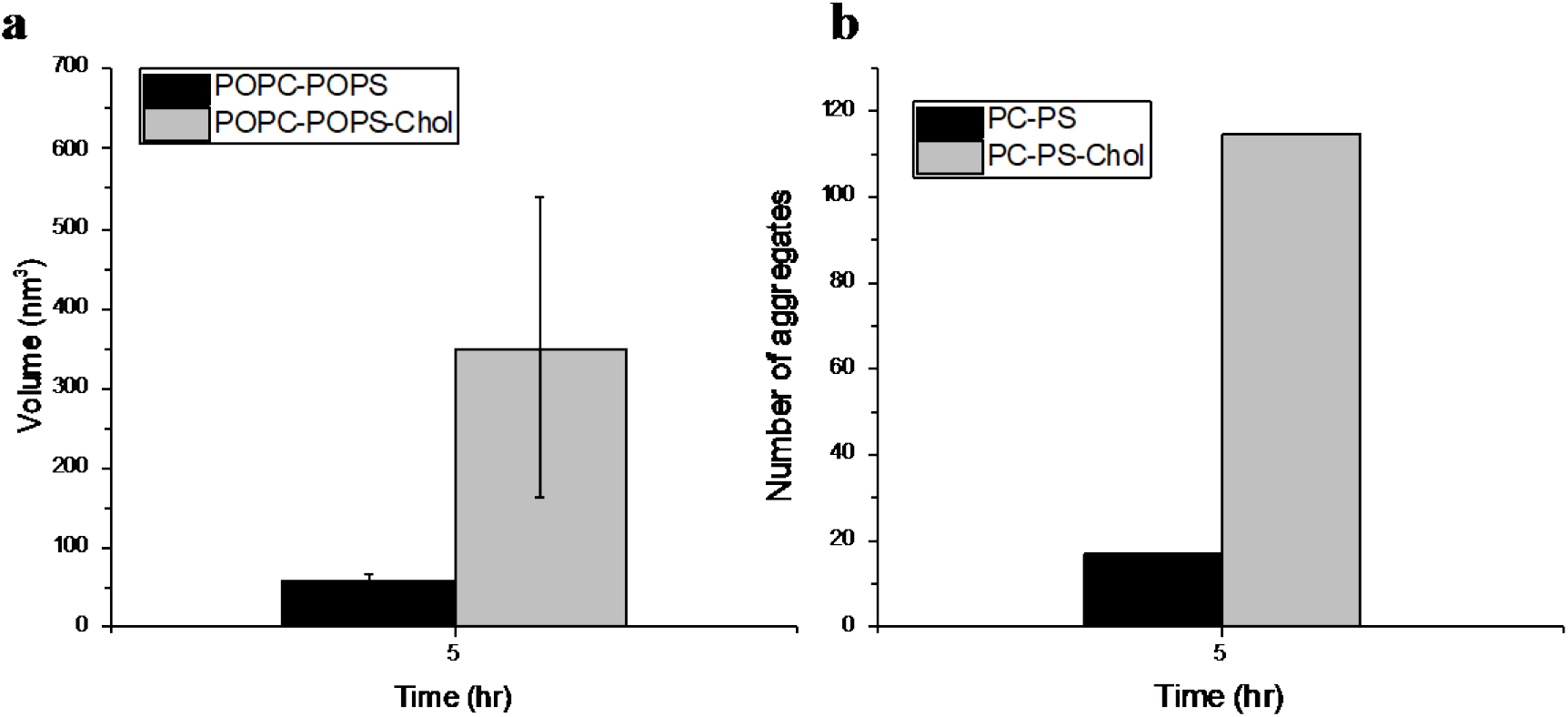
Comparison of aggregation of 10 nM Aβ42 in presence and absence of cholesterol in lipid bilayer. (a) Bar plot compares the volume of aggregates formed after 5 hr of Aβ42 incubation on lipid bilayer with (grey bar) and without (black bar) cholesterol. (b) Bar plot shows the presence of higher number of aggregates on cholesterol containing lipid bilayer (grey bar) after 5 hr of time-lapse experiment.

Next, we used AFM time-lapse imaging to follow the dynamics of aggregates on the lipid bilayer surface (Fig. 3). Along with the formation of the new aggregates, as shown in Fig. 1, some of the aggregates were found to desorb from the surface. The aggregates highlighted in Figs. 3a and 3b with a dotted circle disappear from the surface within 10 min, whereas the neighboring aggregates stay on the surface. The roughness values for an area with no aggregates and the area from which the aggregates dissociated were similar (Fig. 3c). Inspection of both the AFM image and comparison of the roughness value for the area from which the aggregate dissociated reveals that no damage to the bilayer occurred due to the dissociation. Thus, the lipid bilayer is not damaged by the formation or dissociation of large aggregates.

**Figure 3.**
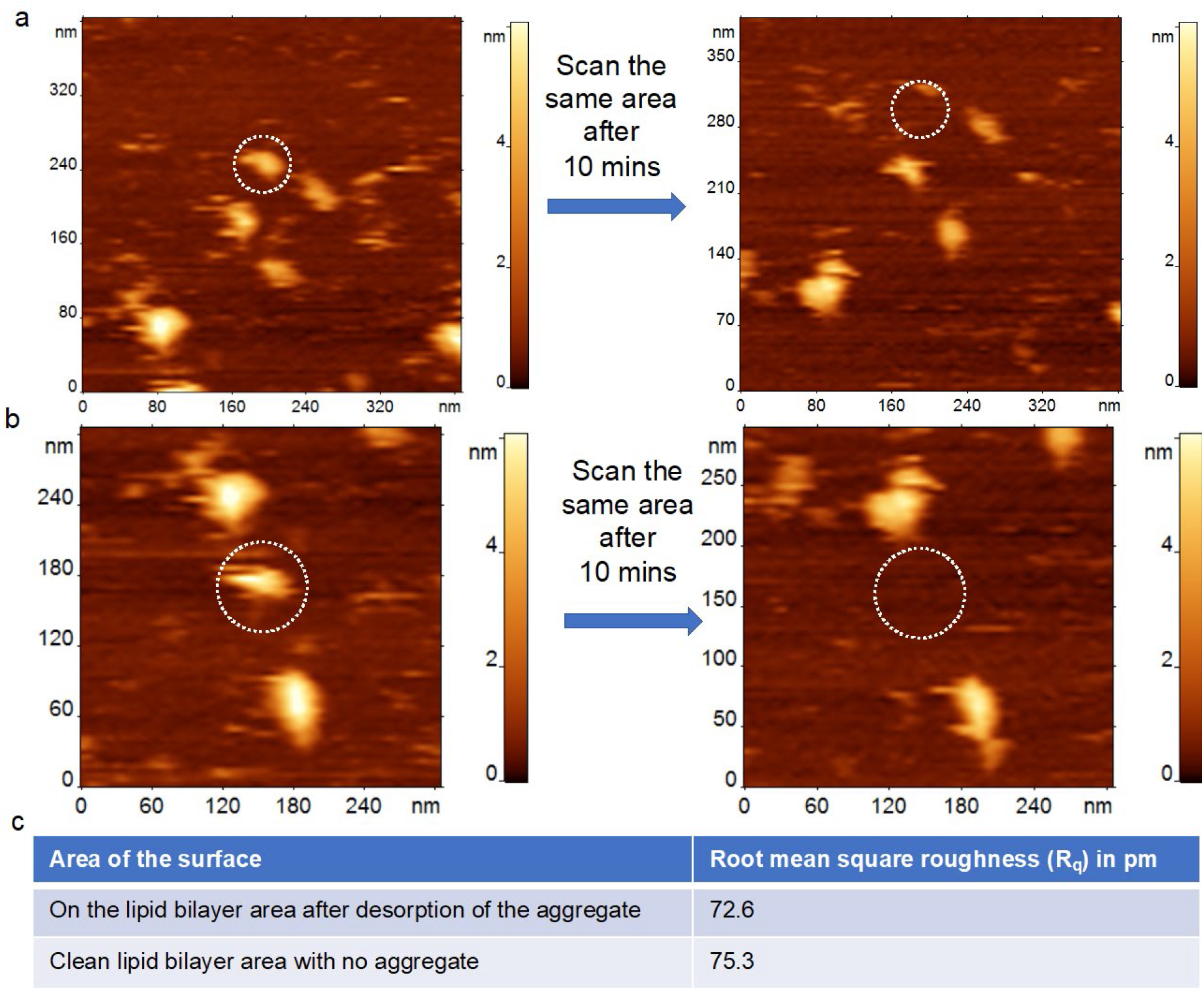
PC-PS-Chol lipid bilayer remains intact after the aggregates leave the surface. **(a-b) The left panel shows the** AFM topographic images of bilayer where one of the aggregates is encircled with dotted line. The right panel shows the same area after 10 min, and the dotted circle indicate the absence of that particular aggregate. (c) The table shows the root mean square roughness values of the particular area after the aggregate leaves the bilayer surface, which is similar to that of a clean bilayer area, indicating the intactness of the bilayer.

To further verify that the aggregates dissociate from the PC-PS-Chol bilayer to the bulk solution and accumulate there, we collected an aliquot from the solution on top of the bilayer surface and deposited it on APS-functionalized mica. Fig. 4a shows the aggregates, after 24 hr of incubation, of 10 nM Aβ42 that accumulated in the bulk solution after dissociation from the lipid bilayer surface. The number of aggregates was then compared to those formed in absence of the bilayer, under the same experimental conditions (Fig. 4b). The grey bars depict the number of aggregates observed in the aliquot of solution from above the bilayer surface, whereas the black bars show the aggregate from the control. These data demonstrate that a large amount of aggregates are released from the PC-PS-Chol bilayer surface into the bulk solution.

**Figure 4.**
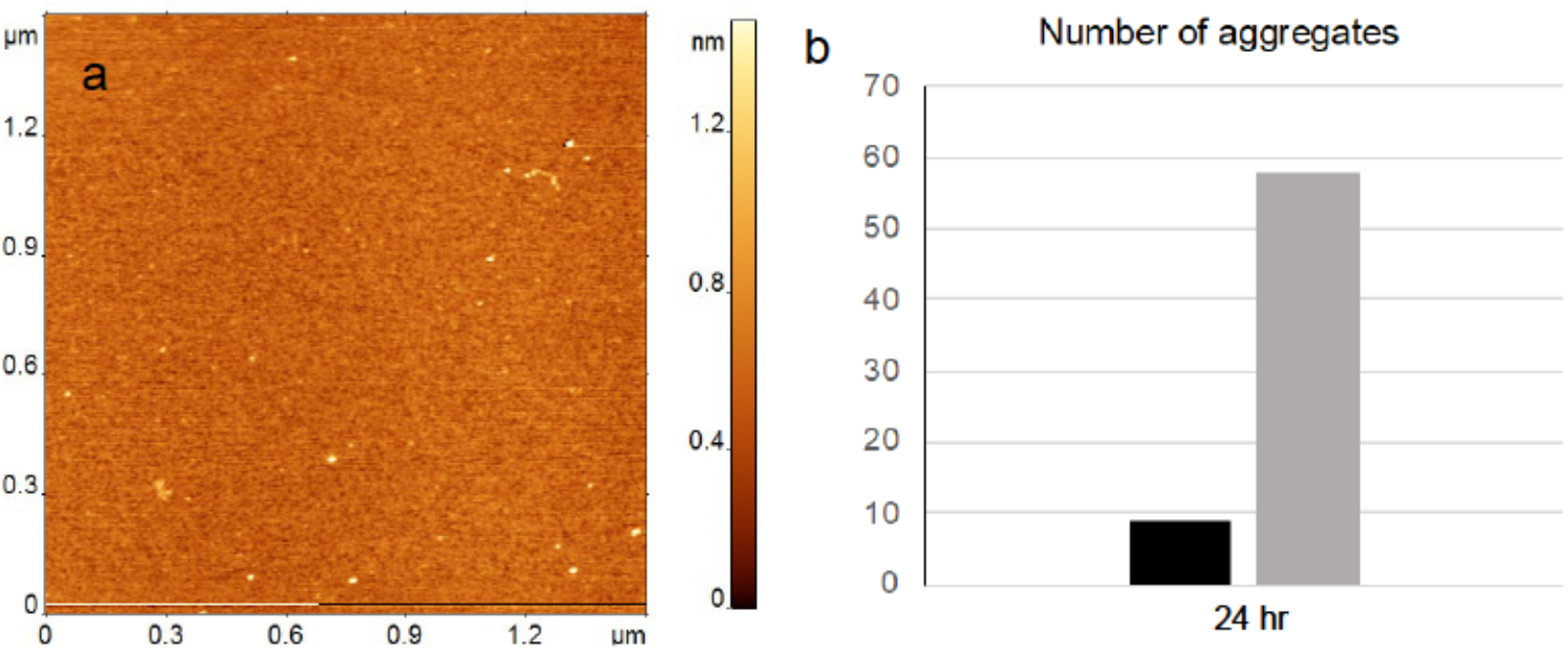
Accumulation of aggregates in the bulk solution after dissociation from the PC-PS-Chol lipid bilayer. (a) AFM image of APS functionalized mica surface after depositing the aliquot obtained from top of the PC-PS-Chol lipid bilayer after 24 hr incubation of 10 nM Aβ42. (b) The bar plot compares the number of aggregates observed in the solution in presence (grey bar) and absence of lipid bilayer (black bar).

### 2. Computer modeling

To elucidate the molecular mechanism behind the enhanced aggregation of Aβ42 on bilayers containing cholesterol, we employed all-atom molecular dynamics (MD) simulations. Briefly, the same two PC-PS bilayers as in experiments, one with and one without cholesterol, were assembled, Aβ42 monomer was placed 5 nm center of mass (CoM) distance from the bilayer core, and the dynamics was recorded for 5 μs.

Snapshots of the Aβ42 monomer in presence of the PC-PS-Chol bilayer at different time points along the simulation trajectory are shown in Fig. 5a. The initial structure (0 ns) is essentially unstructured with two short α-helical regions, shown in blue. After 1.75 μs, three β-sheet segments (shown in yellow) appear, which further grow so that after 3.5 μs two extended β-sheet segments are formed. Thus, interaction of the Aβ42 monomer with the surface induces a conformational change in the monomer that re-arranges the protein backbone. The graph in Fig. 5b summarizes time-dependent overall change of the secondary structure for the Aβ42 monomer. It demonstrates that over time the β-structure content increases, initially fluctuating below 10% (0-3 μs), then jumping to 30% (3-4 μs), before finally stabilizing around 20% (4-5 μs).

**Figure 5.**
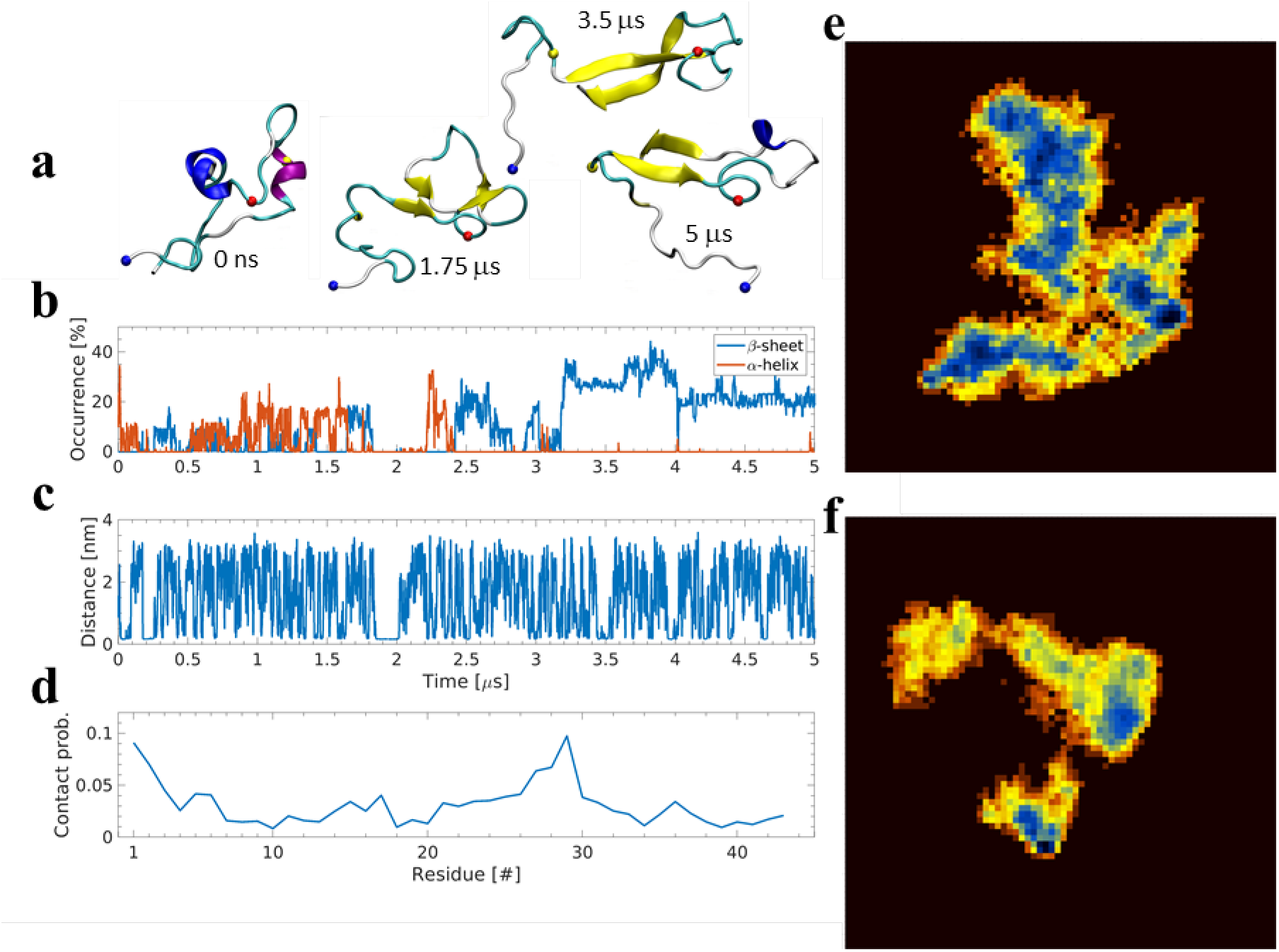
Interaction of Aβ42 monomer with PC-PS-Chol bilayer. **a)** Snapshots of Aβ42 monomer conformations. N-terminus, residue 14, and residue 23 Cα atoms are depicted by blue, yellow, and red spheres, respectively. **b**) Evolution of α-helical, red, and β-structure, blue, secondary structure elements. **c**) Minimum distance between the Aβ42 monomer and the membrane surface. **d**) Probability of interaction between protein residues and the bilayer. **e**) Free energy surface based on dihedral principle component analysis for the monomer on PC-PS-Chol bilayer. Plot shows the first two principle components and the deepest minimum is 4.8 RT. **f**) Free energy surface for the monomer on PC-PS bilayer. The plot uses the first two principle components and the deepest minimum is 6.54 RT. For panels e and f darker blue signifies deeper minima.

Dramatic conformational changes of the Aβ42 monomer do not require strong binding of the molecule to the surface. On the contrary, the interaction is transient, as illustrated in Fig. 5c, which shows the time-dependent change of the distance between the Aβ42 and the lipid bilayer. Additionally, we analyzed the residue-wise probability of interaction of the monomer with the surface. These results are shown in Fig. 5d, which reveal residues 1-7, 15-19 (the central hydrophobic core, CHC), and 21-31 as major segments of the protein responsible for the interaction with the surface.

To further characterize the conformational dynamics of the monomer, we obtained the energy landscape by the use of dihedral principle component (dPC) analysis. The results shown in Fig. 5e demonstrate that the energy landscape is arranged in three trenches of interconnected minima. Each trench contains several low energy minima, the lowest being ~4.8 RT, with numerous other minima within 1 RT (at 300K). A list of a number of minima ranked based on energy is provided in Table S1. The table shows that approximately 10% of the conformations sampled during the simulation have energies within 1RT of the lowest energy minimum.

To evaluate the role of cholesterol on the conformational dynamics of Aβ42 monomer, a similar energy landscape was obtained for the monomer interacting with the bilayer without cholesterol (PC-PS only), shown in Fig. 5f. Although interaction of Aβ42 monomer with this surface also induces the formation of β-structure (Fig. S4a), the energy landscape is dramatically different. The energy landscape is divided into three well separated regions, each with local minima (shown in blue). Furthermore, the absence of cholesterol has led to overall fewer conformations with low barriers, as seen in the decreased number of conformations within 1 RT of the lowest minimum in Table S1. Additionally, the monomer is less likely to interact with the bilayer in the absence of cholesterol, seen as decreased interaction probability in Fig. S4b.

### Modeling of the assembly of Aβ42 dimers on membrane surfaces

Next, we simulated the assembly of the dimer to test the role of cholesterol on this first step of the Aβ42 aggregation process. A free Aβ42 monomer, Mon 2, was placed at 4 nm CoM distance to the bilayer-bound monomer, Mon 1, and the system was simulated for 8.4 μs. As seen in Fig. 6a, after ~260 ns, the CoM value between monomers drops to ~2 nm, suggesting that the dimer is rapidly formed. The assembled dimer is able to dissociate and re-associate with the bilayer during simulation, seen as an increase and decrease in the distance between dimer and the bilayer, Fig. 6b. The dimer stays on the surface for a total accumulated time of ~3320 ns, with the longest continuous residence time being ~1479 ns. As depicted in snapshots on Fig. 6a, the dimer undergoes structural transformation. The added monomer initially has helical structure (I) that is gradually transformed, first by re-arrangement of the termini, from far to near each other (II), then going on to adopt β-structure with terminal anti-parallel β-strands (III), and finally, only having β-strands (IV). This transformation is also observed on the secondary structure plot in Fig. 6b, which shows that, as the simulation progresses, the β-structure content of each monomer gradually increases. For Mon 1, Fig. 6c, the β-structure increases from ~20% to ~40%, then returns to ~20%, before increasing back to ~40% and remaining to the end of the simulation. A similar trend is seen for Mon 2, Fig. 6d, which sees an increase of β-content to ~20% during the simulation. The secondary structure maps on Fig. S5 further reveal the location of β-strands. For Mon 1, residues 16-19, 28-33, and 36-41 adopt β-strand conformation throughout the simulation. Mon 2, on the other hand, adopts β-strands in residues 2-5, 11-14, and 19-21, Fig. S5.

**Figure 6.**
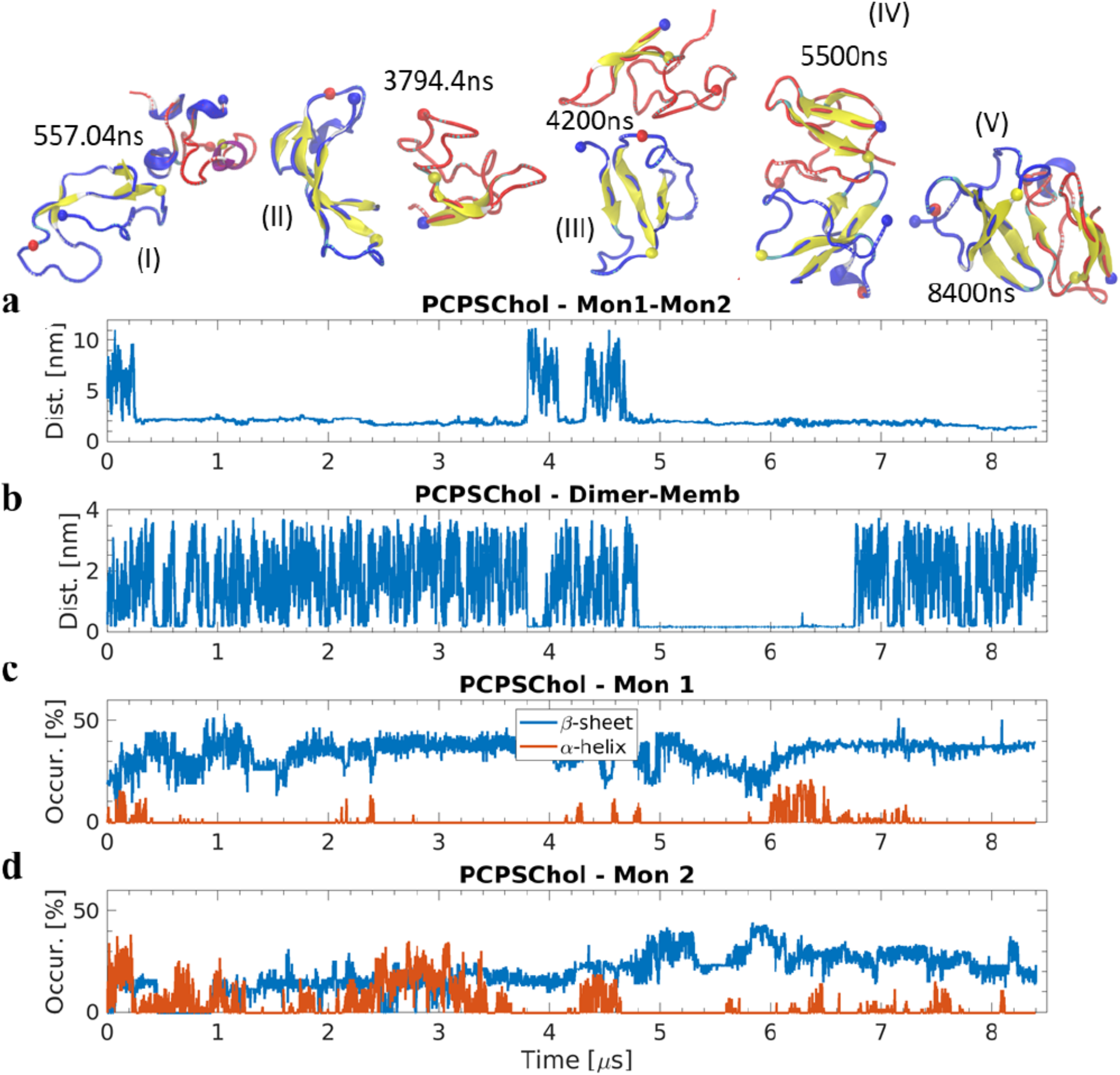
Dimer formation between bilayer-bound Aβ42 monomer and free monomer on PC-PS-Chol bilayer. **a)** Center-of-mass distance between the two monomers. Snapshots depicting the structural transition of the dimer are shown for select timepoints; Mon 1 is depicted in blue, Mon 2 in red, and N-terminus, residue 14, and residue 23 are depicted by blue, yellow, and red spheres, respectively. **b**) Minimum distance between Aβ42 and bilayer surface. **c-d**) Evolution of secondary structure elements within the monomers in presence of PC-PS-Chol bilayer. α-helix is depicted in red and β-sheet in blue.

We then evaluated the effect of cholesterol on the affinity of dimers to the surface by comparing with dimerization on the bilayer without cholesterol. On the PC-PS bilayer, dimerization occurs approximately two times slower, after ~420 ns, compared to the cholesterol bilayer, as seen on Fig. S6a. Furthermore, the total time the dimer spends interacting with the bilayer (a total of ~1631 ns) is ~2 times shorter than on the cholesterol bilayer, and the longest continuous interaction is almost six times shorter, being ~256 ns long, shown in Fig. S6b. Importantly, whereas on the cholesterol bilayer the dimer gained β-structure, in the absence of cholesterol the dimer sees a gradual decrease to ~20%, shown on Figs. S6c-d and the secondary structure map Fig. S7.

Next, we compared the affinity of the dimers to the two bilayers, Fig. S8. The data in Fig. S8a reveal the dimer has significantly higher interaction probability with the cholesterol containing bilayer. The N-terminal residues of both monomers within the dimer are important for the interaction with the bilayer, in particular segments 1-8 and 1-18. However, the affinity for the bilayer without cholesterol shown in Fig. S8b is several times lower, suggesting that cholesterol within the bilayer increases the dimers interaction with the surface. A significant difference is also observed in the geometrical parameters of the two dimers, with the dimer on the cholesterol bilayer experiencing greater backbone RMS fluctuations (up-to 0.5 nm) and deviation (up-to 0.5 nm) compared to the dimer formed on the PC-PS bilayer, depicted in Fig. S8c-d.

## Discussion

Overall, we provided direct evidence of the enhancing effect of cholesterol on the aggregation of Aβ42 in the presence of lipid bilayers. We found that aggregation is a dynamic process in which assembled aggregates of Aβ42 dissociate into the solution without any damage to the bilayer. Computer modeling revealed that Aβ42 monomers dramatically change conformations, facilitating their assembly into dimers. These data further support our model for the membrane mediated catalysis of Aβ42 self-assembly allowing for Aβ42 aggregation at physiologically low concentration, with an additional catalytic effect provided by cholesterol imbedded within the lipid bilayer. Below, we discuss these findings in details as well as the relevance and importance of cholesterol for the development of AD.

### Catalytic effect of cholesterol on Aβ42 aggregation

Time-lapse AFM imaging was a direct and non-ambiguous way to characterize acceleration of Aβ42 aggregation on the membrane surfaces. The number of aggregates grows dramatically faster on the bilayer containing cholesterol, which is graphically shown in Fig. 2. AFM is capable of measuring the size and number of aggregates directly, and it is observed that both the aggregate size and number are larger in the presence of cholesterol-containing membrane. Note that all measurements were made at Aβ42 concentration of 10 nM, which is close to the physiological concentration of Aβ42 [8, 9]. The assembly of aggregates is a dynamic process in which assembled aggregates can dissociate into the solution (Figs. 3 and 4). Importantly, the on-membrane aggregation process is not accompanied by damage to the bilayer (Fig. 3), suggesting that no insertion of the aggregates, required for the pore formation, occurs and the assembly of aggregates is an on-surface process. This observation is in line with paper [23] in which the formation of pores was not observed at physiologically low concentrations of Aβ oligomers. The formation of pores required protein concentrations two orders of magnitude higher than used in our studies.

Overall, the experimental results clearly reveal the key aspects which have not been addressed previously as far as amyloid aggregation in membrane environment is concerned. First, Aβ42 can form aggregates, even in physiologically relevant low protein concentration, and this aggregation is further facilitated by the presence of cholesterol in membranes. The aggregates formed under these conditions do not damage the lipid bilayer when dissociating from the surface. This observation highlights the existence of a possible alternative mechanism, where membrane pores or damage to the lipid bilayer is not the only pathway behind the toxic effects of oligomers.

Given that cholesterol incorporated within the phospholipid bilayer is inside the bilayer and Aβ42 is not inserted into the bilayer, the acceleration effect of cholesterol for the on-surface aggregation process of Aβ42 is not an expected phenomenon. The mechanism has been clarified in the molecular dynamics simulations.

### Computer modeling of interaction of Aβ42 with the cholesterol-bilayer

As is shown in Fig. 5, the interaction of the Aβ42 monomer with the cholesterol-containing bilayer leads to a dramatic change in the protein conformation. The monomer rapidly adopts helical and β-structure conformations before stabilizing in a conformation with anti-parallel β-sheet. Similar conformational change was observed on the bilayer without cholesterol, but the presence of cholesterol is characterized by a greater probability of monomer-bilayer interactions. This has a dramatic effect on the sampling of Aβ42 conformations, clearly seen in Figs. 5e-f, which show the effect of cholesterol on the energy profiles of the monomer in presence of the two bilayers. The energy landscape for Aβ42 interacting with the PC-PS-Chol bilayer is characterized by three connected trenches with local minima. Importantly, these trenches are separated by very low barriers, suggesting an easy conformational transition of Aβ42 within this conformational space. In the absence of cholesterol, three well separated regions are observed (Fig. 5f) and these are separated by higher barriers. Taken together, these results suggest that the bilayer containing cholesterol is able to enhance the structural sampling of the monomer by lowering the barriers between states, and is potentially capable of inducing a state that is more favorable to transition to fibril structure.

The conclusion on the adoption of aggregation-prone conformations by Aβ42 monomers, due to interaction with the surface, is further supported by the simulation of dimer assembly (Fig. 6). In this simulation the Aβ42 monomer approaching from solution to the membrane-bound monomer rapidly forms a dimer, followed by the conformational restructuring of the dimer. A similar process is observed in simulations of the Aβ42 dimerization on PC-PS membrane without cholesterol; however, it occurs two times slower. Furthermore, a dramatic difference in conformation is observed, wherein the dimer formed on the cholesterol-containing bilayer experiences an increase in β-strands, while the dimer on PC-PS loses β-structure (Fig. S5 and S7). Another significant difference is the distribution of the secondary structure elements. The dimer formed on the cholesterol bilayer has large segments with β-strands in both monomers (Mon 1 and Mon 2), while the dimer on the non-cholesterol bilayer only has small β-strands in Mon 1, the initially bilayer-bound monomer, before they shrink and Mon 2 adopts small β-strands. This is also evident from the bilayer residence time, which on the cholesterol-containing bilayer is ~two times longer compared to that on the PC-PS bilayer.

Experimental data show that the on-surface aggregation of Aβ42 occurs without insertion of the protein into the bilayer. The simulation data are in line with this model. Moreover, the interaction of Aβ42 with the surface is transient, and after dissociation from the surface, the protein is able to interact with the surface through a different segment. Simulations also reveal a preference for the monomer to bind the surface with highest probability through the segments between residue 15 and 25 (Fig. 5d). This is also translated to the dimer interactions with the bilayer, with short segments (residues 1-8, 1-18, and 36-40) being more likely to interact with the bilayer, Fig. S8. The Aβ42 dimer is more likely to interact with the cholesterol-containing bilayer compared to the PC-PS bilayer without cholesterol, which translates to a higher continuous interaction time with the bilayer, being approximately six times higher on the cholesterol-containing bilayer.

### Model for the surface catalysis of Aβ42 aggregation and implications to the disease development

According to our model for the aggregation catalysis of amyloid protein by surfaces, the affinity of a protein to the surface is the mechanism explaining surface catalysis of Aβ42 aggregation [13]. As a result, the aggregation was observed at protein concentrations as low as nanomolar. Cholesterol increases the binding affinity of Aβ42 to the PC-PS bilayer, so the elevated catalytic activity of PC-PS-Chol compared with PC-PS is in line with this model. Evidence from literature links AD pathogenesis with an increased concentration of cholesterol [1, 24, 25]. Moreover, cholesterol is normally incorporated into the cellular membranes. Therefore, we hypothesize that one of the mechanisms regulating cholesterol incorporation may disrupt homeostasis, leading to an increase in cholesterol incorporation into the membrane. This model explains the link between the age-dependent increase of the cholesterol concentration in membrane and the AD development.

The formation of pores in membranes is considered as the major neurotoxic effect of Aβ oligomers (reviewed in [26, 27]). However, this model was based on the *in vitro* studies where the Aβ concentration considerably exceeded the physiological concentrations of Aβ, or in experiments with assembly of membranes in the presence of Aβ proteins [28]. According to a recent publication[23], insertion of Aβ42 into membranes with the potential formation of pores requires concentrations of Aβ42 exceeding the physiological ones by several orders of magnitudes. These findings bring concerns to the pore formation model, or the need of additional data to explain such effects of Aβ protein at physiologically relevant concentrations.

Our studies highlight another property of membranes-their catalytic effect, allowing for Aβ42 aggregation at physiologically low concentrations. This provides a novel view on the mechanism of the Aβ42 assembly into aggregates and the role of Aβ42-membrane interactions on the disease development. According to our data, the membrane-assembled Aβ42 aggregates can dissociate from the surface, and may participate in various pathological processes associated with the disease development. Importantly, the majority of aggregates visualized in this work are oligomers, which are considered as the most neurotoxic species among Aβ42 aggregates. Additionally, a previous publication has demonstrated that cholesterol depletion has a neuroprotective effect on amyloid beta toxicity[29]. Therefore, the increase of cholesterol concentration in cell membranes can be a triggering factor for AD, suggesting that controlling the concentration can be a possible venue for prevention of the disease. It has been reported that in addition to cholesterol, lipids, such as sphingomyelin, and glycosphingolipids, such as gangliosides, can contribute to Aβ42 aggregation[30–32]. The elucidation of the role of these lipids in the Aβ42 aggregation is our long-term goal.

## Materials and methods

### Materials

All lipids were purchased from Avanti Polar Lipids, Inc, Alabama, US. Aβ42 was purchased from AnaSpec, Fremont, CA. Chloroform was procured from Sigma Aldrich Inc. The buffer solution that was used in this study is 20 mM HEPES, 150 mM NaCl, 10 mM CaCl2, pH 7.4.

#### Preparation of Aβ42 protein solution

Stock protein solution was prepared as described in the previous publication[10]. Briefly, Aβ42 was dissolved in 1,1,1,3,3,3-hexafluoroisopropanol (HFIP) to break any preformed aggregates. Then HFIP was evaporated by vaccufuge and then the stock solution is prepared in DMSO. Detailed methods is presented in supporting information.

#### Preparation of supported lipid bilayer

To prepare the supported lipid bilayers on mica surface we adopted the protocol described in our previous studies[11, 12]. Detailed method is available in supporting information.

### Computational methods

#### Bilayer assembly

We employed CHARMM-GUI [33] to generate the initial bilayers of POPC:POPS, and POPC:POPS with cholesterol. Bilayers were then energy minimized, heated, relaxed using NVT MD, and run for 150 ns as NPT MD to fully relax the bilayer. Simulations were performed using the Amber16 package [34]. Detailed explanation is given in supporting information.

#### Interaction of Aβ42 with bilayers

Aβ42 monomer (conformation taken from ref. [35]) was added to each bilayer system and, after a short relaxation simulation, run on the special purpose supercomputer Anton 2 for long production runs. Detailed methods provided in supporting information.

#### Interaction between membrane-bound and free Aβ42

A free Aβ42 monomer (from ref. [35]) was added to the last membrane-bound conformation of the previous simulation systems and, after preparatory simulation steps, run on Anton 2 special purpose supercomputer. Detailed description is available in the supporting information.

#### Analysis of MD trajectories

Gromacs suite of programs (v2020) [36], Carma [37], and VMD [38] were used to analyze the obtained simulation trajectories.

## Acknowledgments

This research was funded by National Institutes of Health, grants GM096039 and GM118006 to Y.L.L. Anton 2 computer time was provided by the Pittsburgh Supercomputing Center (PSC) through Grant R01GM116961 from the National Institutes of Health. The Anton 2 machine at PSC was generously made available by D.E. Shaw Research. This work was completed utilizing the Holland Computing Center of the University of Nebraska, which receives support from the Nebraska Research Initiative. Y.L.L, S.B, M.H designed the project. S.B performed the AFM experiments. M.H performed and analyzed the molecular dynamics simulations. All authors wrote and edited the manuscript. Authors thank Thomas D Stormberg for proof reading.

