## supplement for "Cholesterol in membranes facilitates aggregation of amyloid β protein at physiologically low concentrations"

This PDF file includes:

Supplementary Methods
Supplementary Table S1, Figures S1 to S8
Supplementary references

**Supplementary Methods:**

*Preparation of Supported lipid bilayer:*

1-palmitoyl-2-oleoyl-glycero-3-phosphocholine (POPC, PC), 1-palmitoyl-2-oleoyl-sn-glycero-3-phospho-L-serine (POPS, PS) and cholesterol (Chol) were used to prepare the supported lipid bilayers. Briefly, vesicles are deposited onto cleaved mica surface, which was glued onto a glass slide, and kept at 60°C for 1 hr. Following which, the surface was allowed to return to the room temperature. Then the surfaced was rinsed, to remove excess vesicles, and a buffer containing 20 mM HEPES, 150 mM NaCl, pH 7.4 was added onto the formed lipid bilayer. The prepared lipid bilayer was imaged immediately using AFM.

*Preparation of Aβ42 protein solution:*

Preparation of Aβ42 stock solution was similar to the method described in our previous publications [1, 2]. Lyophilized Aβ42 powder (Anaspec, Inc.) was dissolved in 100 μL of 1,1,1,3,3,3-hexafluoroisopropanol (HFIP, Sigma-Aldrich Inc.) at room temperature with sonication. HFIP was then completely evaporated in a vacufuge. Anhydrous DMSO was then added to make the stock solution, which is then kept at -20°C. Working protein solution is prepared by diluting the stock into the imaging buffer.

*Time-lapse AFM imaging:*

AFM imaging, in buffer medium, was carried out in tapping mode using an MFP-3D instrument (Asylum Research, Santa Barbara, CA). The cantilever "E" of MSNL probes (Bruker) were utilized for all imaging purposes. The nominal frequency of the cantilever in buffer was 7-9 kHz with a spring constant of ~0.1 N/m. Scan speed was typically between 1 to 2 Hz.

All lipid bilayers were imaged to check that the surfaces were smooth, clean, and devoid of any unruptured vesicles. Then time-lapse imaging was carried out in the same area of the bilayer after the introduction of 10 nM Aβ42 solution. Image acquisition was stopped after recording each frame to ensure that no damage to the lipid bilayer surface occurred due to scanning.

*AFM data analysis*

The presented AFM images are with minimal processing. The images were flattened (fitted with 1$^{st}$ order polynomial) with FemtoScan software (Advanced Technologies Center, Moscow, Russia). The volume of the oligomers was measured using the grain analysis tool in the software. Then the volume data were plotted as histograms using Origin Pro software (OriginLab, Northampton, MA, USA) and fitted with Gaussian distribution. The mean value of the oligomer volume for each time point was determined using the peak value of the distribution and the error bars represent the standard deviation.

<u>Computational Methods</u>

*Bilayer assembly*

We employed CHARMM-GUI [3] to generate the initial bilayers of POPC:POPS, and POPC:POPS with cholesterol. Each bilayer consisted of 512 lipid molecules, 40 TIP3P waters [4] per lipid, and 102 cholesterol (only for PC-PS-Chol system). Lipid systems were neutralized and kept at 150 mM ionic concentration using NaCl counterions, and converted to AMBER format using the lipid17 force field (an extension and refinement of lipid14 [5]). Bilayers were then energy minimized, heated, and run for 500 ps as NVT ensemble. After which 150 ns NPT MD simulations were performed using a 2 fs integration time step. The simulations employed periodic boundary conditions with a semi-isotropic pressure coupling at 1 bar, a constant temperature of 300 K, non-bonded interactions truncated at 10 Å, and electrostatic interactions treated using particle-mesh Ewald [6]. Simulations were performed using the Amber16 package [7].

*Interaction of Aβ42 with bilayers.*

To investigate the interaction of Aβ42 monomer with the bilayers, we extracted the bilayers from the final frame of the bilayer simulations, added Aβ42 monomer (conformation taken from ref. [8]) at 5 nm center-of-mass (CoM) from the bilayer center, solvated in TIP3P water (in an orthorhombic box [a=b≠c] with side:height ratio 0.75), neutralized with NaCl counter ions, and maintained a final NaCl concentration of 150 mM. Proteins were described using the Amber ff99SB-ILDN force field [9]. Each system then underwent H-mass repartitioning to increase H mass to 3.024 Da allowing for 4 fs time steps [10], following which the systems were energy minimized, heated, run for 500 ps as NVT ensemble, and simulated as an NPT ensemble for 10 ns (using the same parameters as previous bilayer simulations) before being submitted to the special purpose supercomputer Anton 2 for long production runs. Simulations on Anton 2 employed the multigrator algorithm and treated electrostatics using the Gaussian split Ewald method.

*Interaction between membrane-bound and free Aβ42.*

To investigate the interaction between membrane-bound and free Aβ42 species, we used the last membrane-bound conformation of the previous simulation systems and added a monomer (from ref. [8]) to the simulation systems following the same procedure as used initially to add Aβ42 molecules to the bilayer systems. Newly added monomers were placed at 4 nm CoM with respect to the membrane-bound molecules and at 5 nM distance to the membrane core. Simulation parameters and steps were the same as the initial Aβ42-bilayer simulations.

*Analysis of MD trajectories*

Gromacs suite of programs (v2020) [11], Carma [12], and VMD [13] were used to analyze the obtained simulation trajectories.

**Supplementary Table and Figures:**

| Energy [RT] | PC-PS-Chol | PC-PS | Difference [%] |
|---|---|---|---|
| < -1 | 24.27 % | 14.27 % | 70 |
| < -2 | 18.02 % | 10.98 % | 64 |
| < -3 | 9.63 % | 6.94 % | 38 |
| < -4 | 0.98 % | 2.26 % | - 56 |
| < -5 | - | 0.38 % | - |
| < -6 | - | 0.06% | - |

**Table S1. Interaction of Aβ42 monomer with PC-PS and PC-PS-Chol bilayers.** The table shows total number of conformations identified through dihedral principle component analysis that have a particular free energy in units of RT (at 300K).

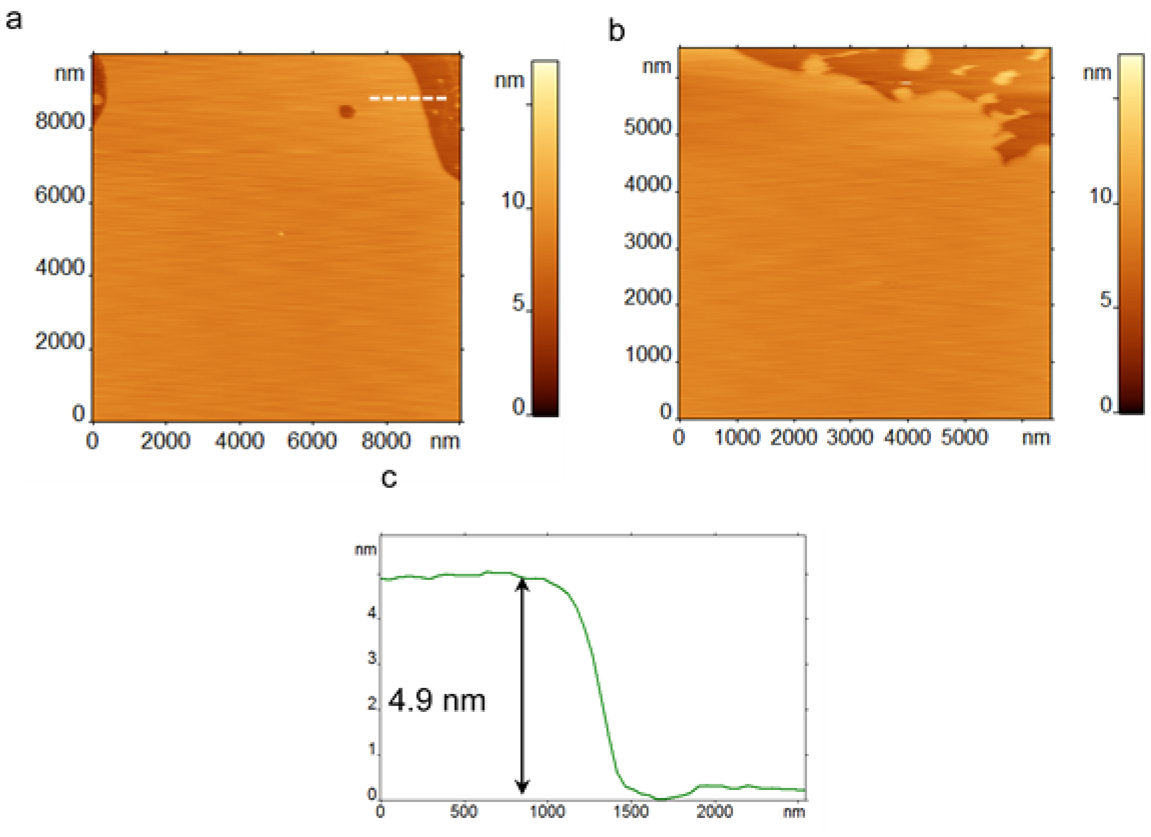


**Figure S1. Homogeneity and smoothness of PC-PS-Chol.** (**a-b**) AFM topographic images of PC-PS-Chol lipid bilayer on two different areas of the sample. Both areas are smooth, homogeneous, and free of any trapped vesicles. (**c**) Cross-section profile of the lipid bilayer. The height value of 4.9 nm indicates the formation of lipid bilayer.

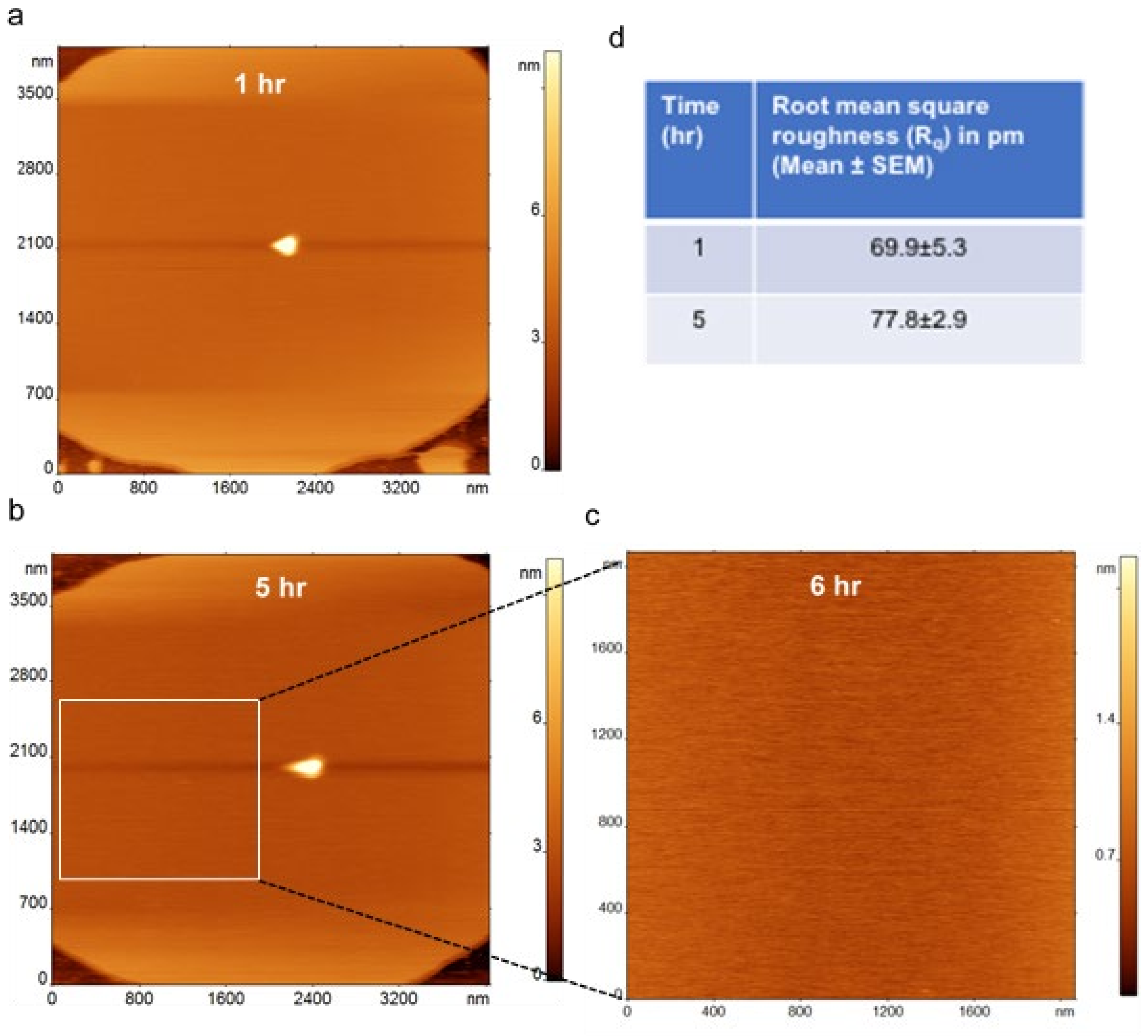


| Time (hr) | Root mean square roughness ($R_q$) in pm (Mean ± SEM) |
|---|---|
| 1 | 69.9±5.3 |
| 5 | 77.8±2.9 |

**Figure S2. Stability of PC-PS-Chol.** AFM topographic images of the same area on the PC-PS-Chol bilayer surface after (**a**) 1 hr and (**b**) 5 hr. (**c**) A zoomed image of the bilayer surface. The bilayer remains clean throughout the experiment and no aggregate like feature was observed. (**d**) The table shows that the roughness of the bilayer remains similar over the time, indicating the smoothness of the bilayer is maintained over the time course.

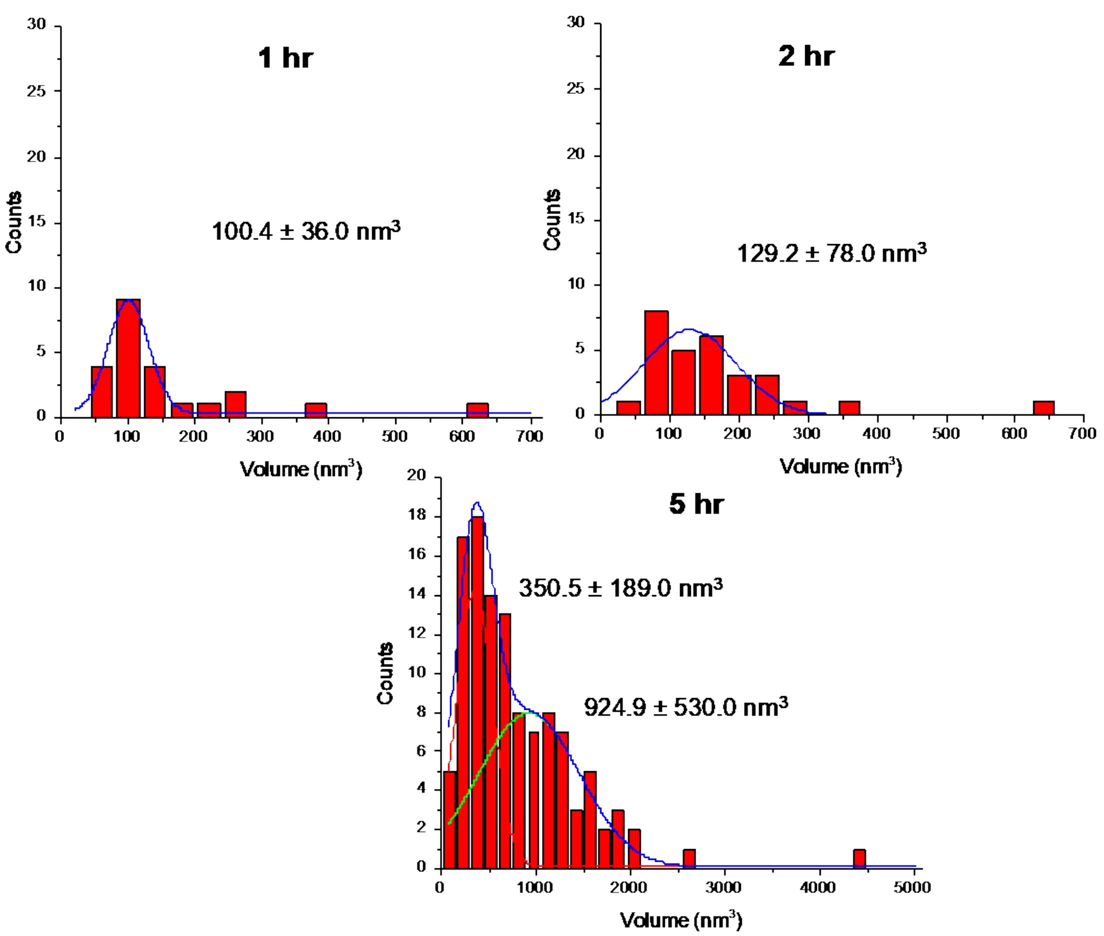


**Figure S3. Quantitative analysis of aggregation on PC-PS-Chol bilayer.** The distribution of the volume of the aggregates appeared on the bilayer surface at 1 hr, 2 hr, and 5 hr time-points.

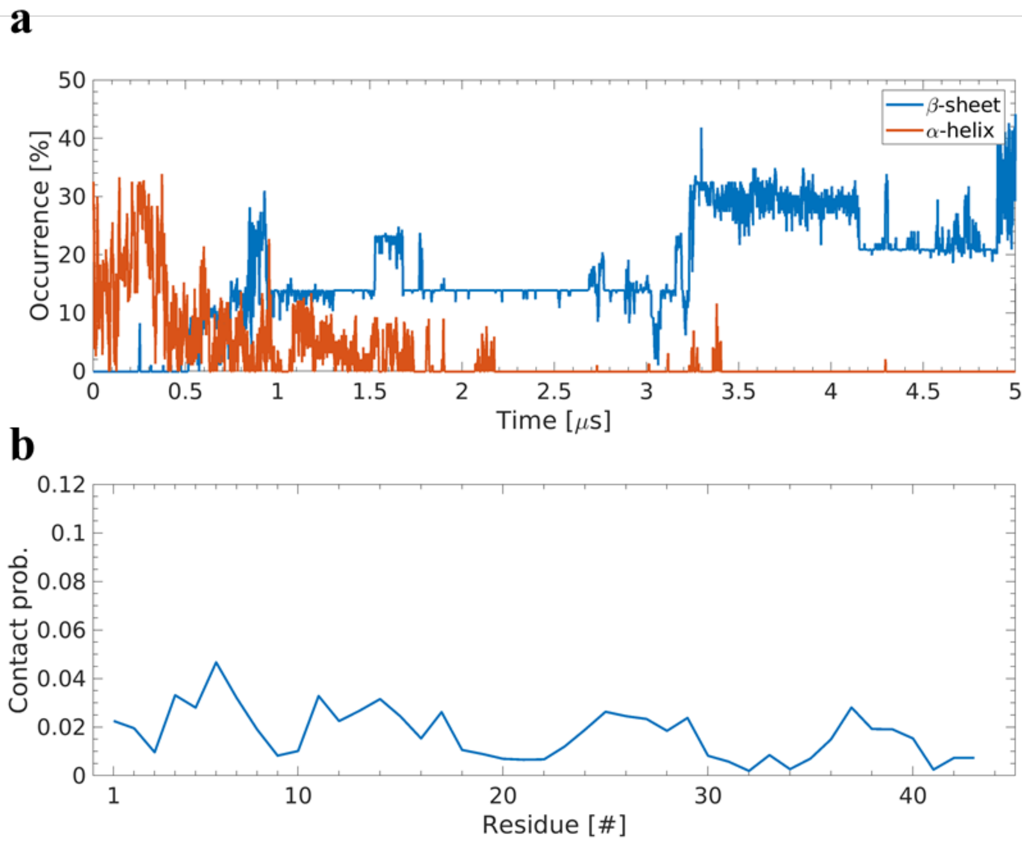


**Figure S4. Interaction of Aβ42 monomer with PC-PS bilayer. a**) Secondary structure analysis showing the evolution of α-helical and β-structure for monomer in presence of PC-PS bilayer. **b**) Probability of interaction between Aβ42 monomer residues and the bilayer.

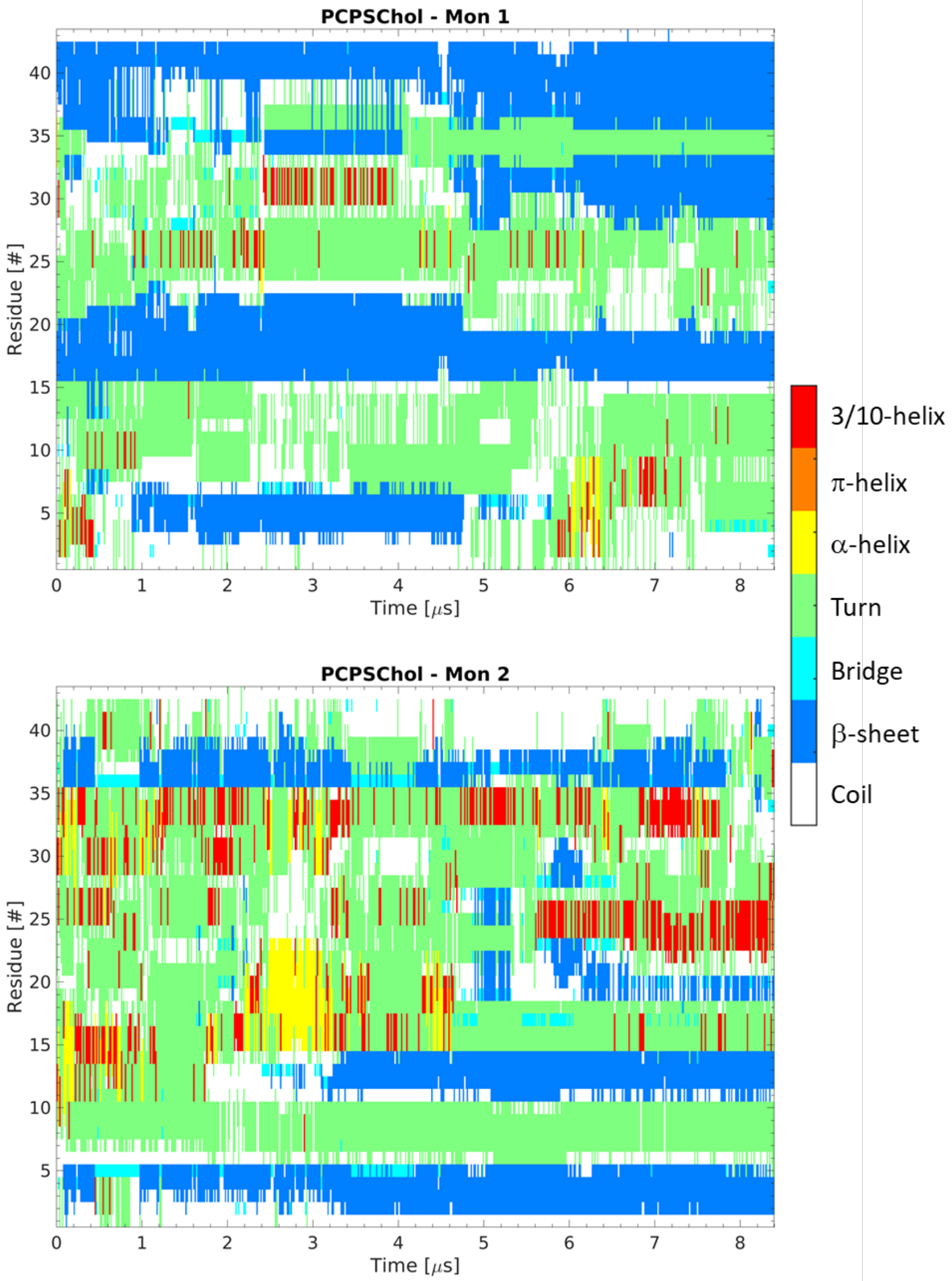


**Figure S5. Secondary structure map of the evolution of structure within the monomers of the dimer interacting with PC-PS-Chol bilayer.** Secondary structure for initial membrane-bound monomer (Mon 1), top, and the initially free monomer (Mon 2), bottom, were determined using STRIDE through VMD.

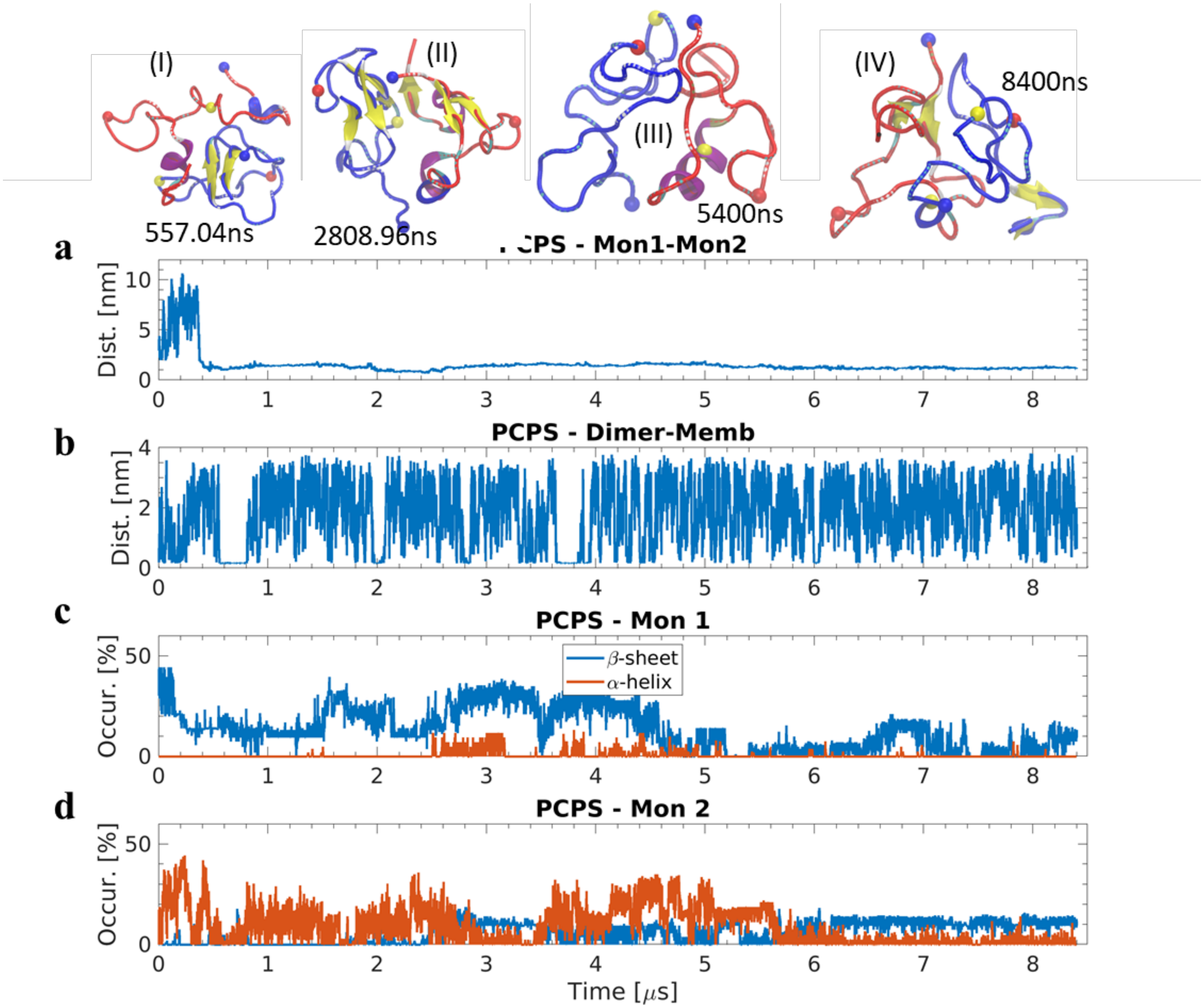


**Figure S6. Dimer formation between bilayer-bound Aβ42 monomer and free monomer on PC-PS bilayer. a**) Center-of-mass distance between the two monomers. Snapshots depicting the structural transition of the dimer throughout the simulation are shown with following highlights: Mon 1 is depicted in blue, Mon 2 in red, and N-terminus, residue 14, and residue 23 are depicted by blue, yellow, and red spheres, respectively. **b**) Minimum distance between Aβ42 and the bilayer surface. **c-d**) Evolution of secondary structure for each monomer versus simulation time on PC-PS.

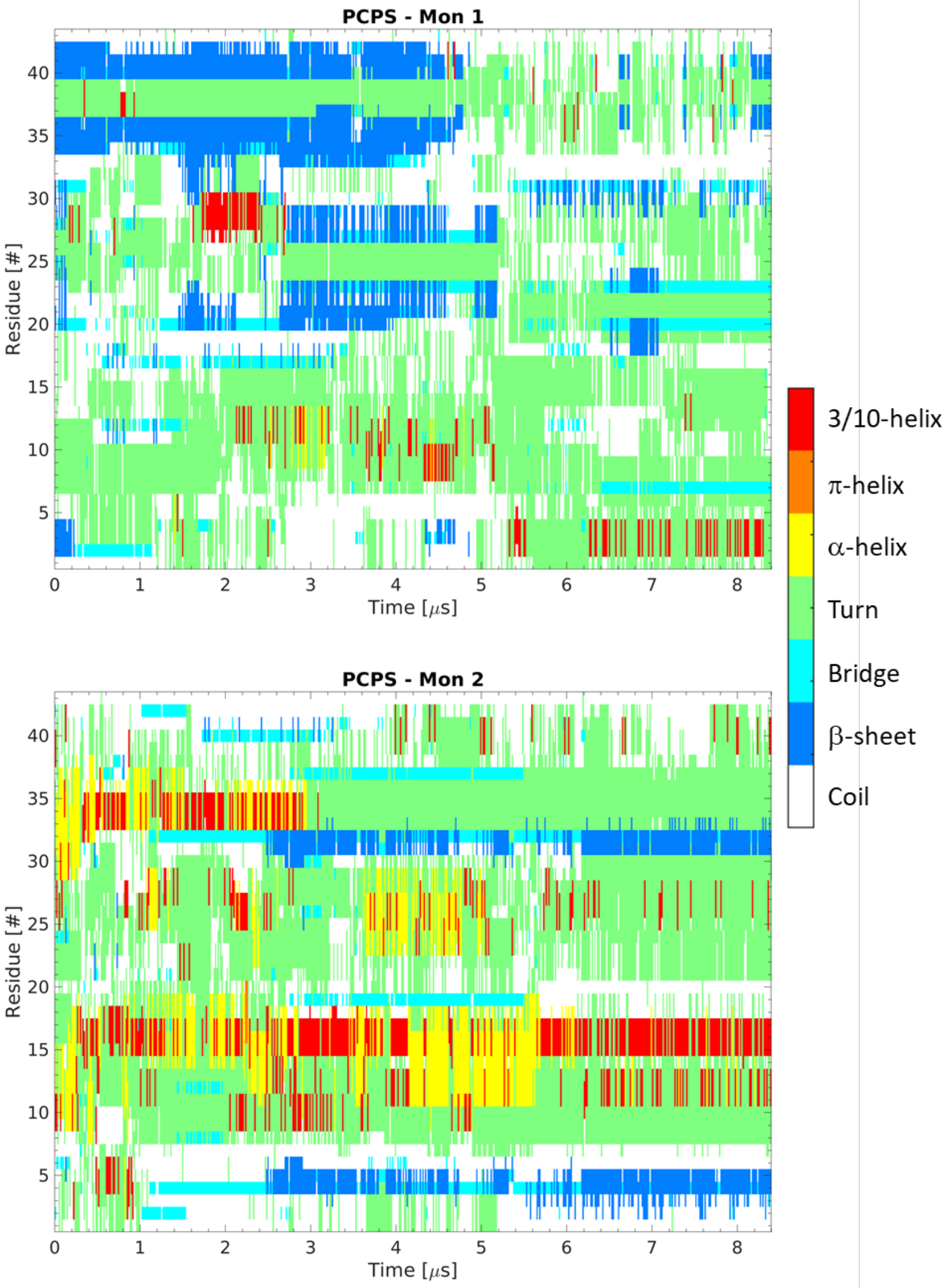


**Figure S7. Secondary structure map of the evolution of structure within the monomers of the dimer interacting with PC-PS bilayer.** Secondary structure for initial membrane-bound monomer (Mon 1), top, and the initially free monomer (Mon 2), bottom, were determined using STRIDE through VMD.

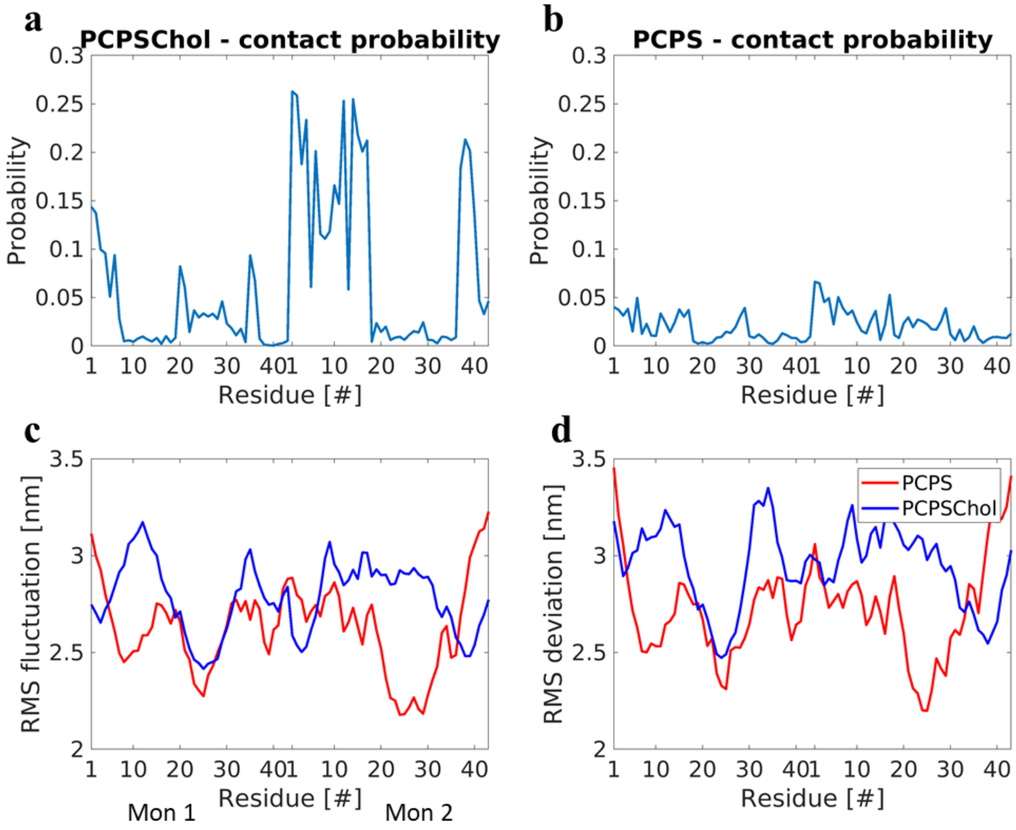


**Figure S8. Dimerization of Aβ42 on PC-PS and PC-PS-Chol bilayers.** Probability of interaction of Aβ42 monomer residues with the, **a**) PC-PS-Chol and **b**) PC-PS bilayers. Mon 1, the monomer initially bound to membrane surface, is represented by the values on the left in the graphs. **c**) Average per-residue root mean square fluctuation of Cα atoms compared to the initial monomer structure. Blue graph depicts the dimer from the PC-PS-Chol bilayer, while red is the dimer on PC-PS. **d**) Average root mean square deviation in the Cα atom positions compared to the initial monomer structure. Color coding is same as panel c.
